# MorphData: Automating the data extraction process of morphological features of microglial cells in ImageJ

**DOI:** 10.1101/2021.08.05.455282

**Authors:** Ana Bela Campos, Sara Duarte-Silva, António Francisco Ambrósio, Patrícia Maciel, Bruno Fernandes

## Abstract

Microglial cells are the first line of defense within the central nervous system, with morphological characterization being widely used to define their activation status. Most methods to evaluate microglia status are manual, and, therefore, often biased, inaccurate, and time consuming. In fact, the process to collect morphological data starts with the acquisition of photomicrographs from where images of single cells are extracted. Then, the researcher collects the morphological features that characterize each cell. However, a manual data collection process from single cells can take weeks to complete. This work describes an open-source ImageJ plugin, *MorphData*, which automatizes the data extraction process of morphological features of single microglial cells. The plugin collects, processes, and organizes features associated with cell complexity and ramification. In a computer with limited computing power, *MorphData* was able to handle 699 single cells in less than 14 minutes. The same process, if performed manually, would take almost 19 working days. Overall, *MorphData* significantly reduces the time taken to collect morphological data from microglial cells, which can then be used to study, understand, and characterize microglia behavior in the brain of human patients or of animal models of neurological and psychiatric diseases.

## 1. Introduction

Microglial cells represent a population of macrophages-like cells in the central nervous system, with a broad range of roles in neurodevelopment, synaptic plasticity, and brain protection and repair ^[1]^. Hence, the morphological characterization of these cells is of the utmost importance to ascertain and establish their state in particular conditions. It is known that microglia morphology and function are closely related ^[2]^. In fact, in response to injury, microglial cells undergo morphological and functional changes, changing from a highly ramified into an amoeboid-like shape ^[3]^. This implies that a rigorous analysis of microglia morphology data is of essence for the understanding of cellular behavior ^[3, 4, 5]^.

The collection of morphological data goes through several steps. First, one is required to obtain photomicrographs from where images of cells can be extracted. Then, ImageJ is required for image processing ^[6]^. Being an open-source software, it is frequent to find macros and plugins, conceived by the community, that provide ImageJ with extra features. Examples include *SlideJ* ^[7]^, *aNMJ-morph* ^[8]^, and *ImageSURF* ^[9]^, and *Simple RGC* ^[10]^, among others. A different example is provided by Heindl and colleagues ^[11]^, where the authors propose a morphological analysis method outside ImageJ, using a closed and proprietary programming language and numeric computing environment.

For the morphological analysis of microglial cells, ImageJ provides two key plugins: (i) *AnalyzeSkeleton (2D/3D)* ^[12]^, which tags skeletal features relevant to cell ramification, and (ii) *FracLac* ^[13]^, which quantifies cell surface and size, soma thickness, and the cylindrical shape of cells. The use of both plugins is recommended, as cell ramification data are complementary to cell complexity ^[14]^. However, while the latter is applied over single cell images, the former is typically applied to entire photomicrographs, thus producing results with significant noise. Hence, we aimed to develop a protocol that allowed the application of both plugins over single cells, not only reducing the amount of noise that comes from analyzing larger and noisy photomicrographs, but also solving the problem of stacked cells. This comes, however, with a significant time cost when collecting the morphological features that characterize each cell. In fact, the process to obtain such morphological features, when performed manually over each cell, is a demanding, repetitive, and laborious task, that can take several weeks to complete. Another potential issue is the human error associated with the data collection process. This sets the need for the *MorphData* plugin.

This manuscript describes the design, implementation, and use of a new plugin that automatically runs and collects morphological features for single cells in a matter of minutes, significantly reducing the time spent on the data collection process. The goal of *MorphData* is set on optimizing the data collection process of morphological features.

## 2. Materials and methods

### 2.1 Ethics statement

All procedures with mice were conducted in accordance with the ARRIVE 2.0 guidelines (Animal Research: Reporting *In Vivo* Experiments). Animal facilities and the people that worked directly in animal procedures were certified by the Portuguese regulatory – *Direção Geral de Alimentação e Veterinária*, license number 020317. All animal procedures were approved by the Animal Ethics Committee of the Life and Health Sciences Research Institute, University of Minho (SECVS 120/2014), and conducted in consonance with the European Union Directive 2010/63/EU. Health monitoring was performed according to the Federation of European Laboratory Animal Science Associations guidelines, where the Specified Pathogen Free health status was confirmed by sentinel mice maintained in the same animal housing room.

### 2.2 Animal maintenance

Two groups, control (CTR) and experimental group (EX) mice on a C57BL/6J background, were considered. Animals were maintained in a conventional animal facility and under standard laboratory conditions, which includes an artificial 12h light/dark cycle, lights on from 8:00 am to 8:00 pm, an ambient temperature of 21 ± 1ºC and relative humidity of 50–60%.

### 2.3 Immunofluorescence staining

CTR (n=4) and EX (n=4) mice were deeply anesthetized with a mixture of ketamine hydrochloride (150mg/kg) and medetomidine (0.3mg/kg), and transcardially perfused with phosphate-buffered saline (PBS) followed by 4% paraformaldehyde (PFA) solution (PFA, 0.1 M, pH 7.4, in PBS). Brains were removed and immersed in 4% PFA (48h), followed by 1 week in a 30% sucrose PBS buffer (at 4ºC). Coronal sections were obtained using a vibratome (VT1000S, Leica, Germany) with 40μm of thickness. For staining, the permeabilization in the free-floating sections was performed with PBS-T 0.3% (0.3% triton X-100, Sigma Aldrich, in PBS) for 10 min, followed by immersing the slices in pre-heated citrate buffer (10 mM, pH 6.0; Sigma Aldrich) during 20 min using a thermoblock (D1200, LabNet) set at 80ºC. Once cooled, slices were blocked with goat serum blocking buffer (10% normal goat serum, 0.3% triton X-100, in PBS) at room temperature (RT) for 90 min. After this, the sections were incubated with the primary antibody anti-ionized calcium binding adaptor molecule 1 (rabbit polyclonal IgG anti-Iba-1, 1:600; Wako) overnight at 4ºC. In the next day, sections were incubated with a secondary antibody (Alexa Fluor 594 goat anti-rabbit, 1:1000; ThermoFisher Scientific) during 90 min at RT, protected from light, followed with 4’,6-Diamidin-2-phenylindol (DAPI, 1:1000; Invitrogen) for nuclei staining. Sections were mounted on microscope slides (Menzel-Glaser Superfrost^©^Plus, Thermo Fisher Scientific) and covered with a coverslip (Menzel-Glaser 24–60mm, Wagner und Munz) using aqueous mounting medium (Fluoromount TM, Sigma-Aldrich).

### 2.4 Image acquisition and preparation

Images were acquired using the Olympus Confocal FV1000 laser scanning microscope, with a resolution of 1024×1024px, using a 40× objective (UPlanSApo, N.A. 0.90; dry; field size 624.39×624.39μm; 0.31μm/px), being used to obtain Z-stacked images, which include two distinct channels (red, Iba-1; blue, DAPI). The acquisition settings were the following: scanning speed = 4μm/px; pinhole aperture = 110μm; Iba-1, excitation = 559nm, emission = 618nm; DAPI, excitation = 405nm, emission = 461nm; in a 3-dimensional scenario (X, Y, and Z axis). Four coronal brain sections per animal were imaged in both hemispheres, for a particular region of interest, to yield 4-6 digital photomicrographs per section containing the region of analysis.

The Z-stacked 3D volume images from sections of the region of interest were prepared for microglial morphology analysis using a semi-automatic method, adapted from ^[14]^, to obtain both skeleton and fractal data. However, contrary to the cited method, we went further and obtained binary (white cells on black background) single cells (one cell per file in TIFF format) to feed both the *AnalyzeSkeleton (2D/3D)* and the *FracLac* plugins.

Briefly, after stacking the 3D volume images, the double-color image was split to obtain the Iba-1 label in the red channel, which accurately mirrors the cell profile. Brightness and contrast of the red-channel were adjusted as needed and an unsharp mask was applied. Then, a despeckle filter was used to remove salt and pepper noise, with the threshold option being used and adjusted, as needed, to convert the image into a binary one. Noise was subsequently reduced using despeckle and by removing outliers. After that, random cells from both the original and the binary images were selected with the rectangle tool, using the region of interest to set the same rectangle dimensions for all the selected cells (field size 296×264). Then, after selecting the cells, the paintbrush tool was used to complete and draw the morphology of the cells (always comparing with the original image) and to clean extra signal that is not related to these cells, thus producing a single-cell image without any noise. This process is summarized in Figure 1a and 1b. Sample data are available at *MorphData*’s code repository (github.com/anabelacampos/MorphData).

**Figure 1.**
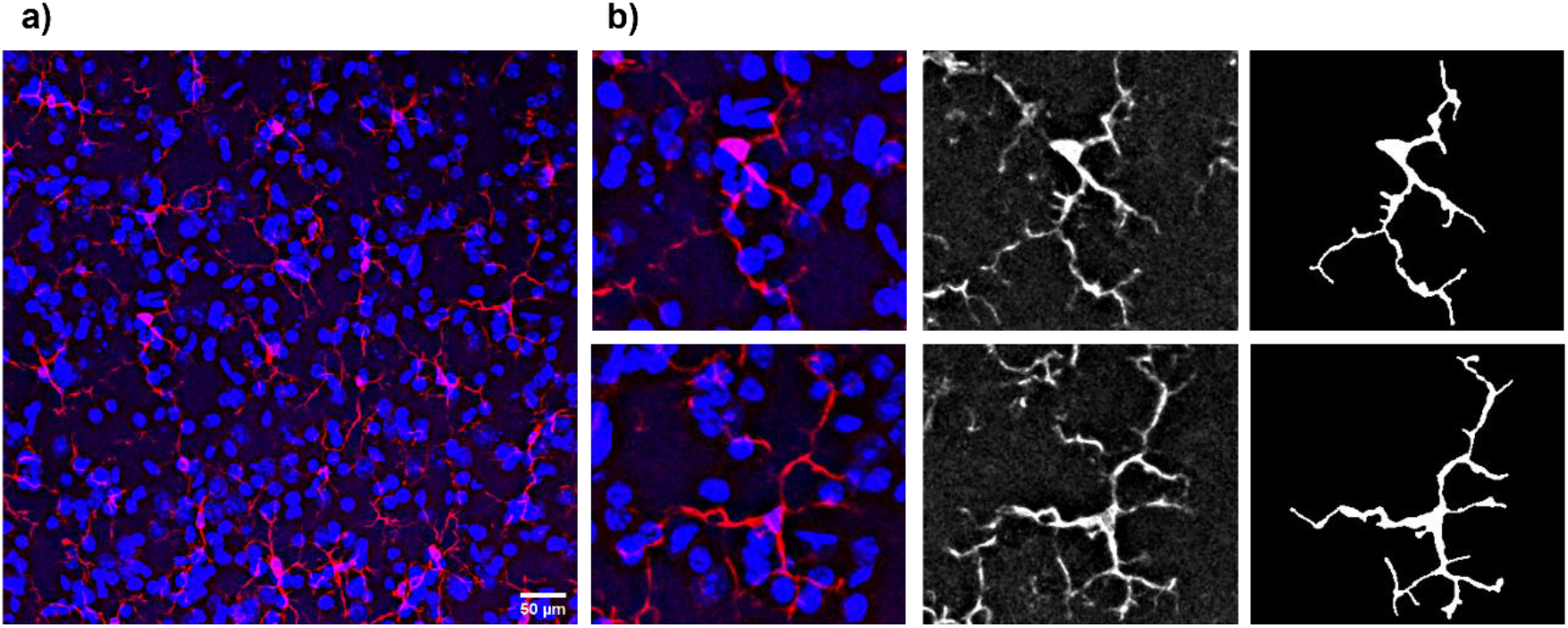
Representative photomicrograph from the region of interest, showing microglial cells (red) and the representation of two binary single microglial cells (gray) from a control mouse. **a)** Original Z-stacked 3D volume photomicrograph. **b)** Going from a noisy cell to a binary single cell (from the left to the right).

699 single-cell images, for both CTR (310 single cells) and EX (389 single cells) groups, were obtained and stored in the file system, in the TIFF format. At this point, the researcher is ready to start collecting the morphological features that characterize each microglial cell using the *MorphData* plugin.

## 3. MorphData architecture and implementation

The *MorphData* plugin was developed using ImageJ Macro language (IJM), a scripting language that allows a developer to control many features of ImageJ. Plugins written in IJM can be programmed to perform sequences of actions, thus automating repetitive processes. It has a set of basic structures, including variables, conditional statements, loops, and user-defined functions. Importantly, IJM allows the developer to access ImageJ functions that are available from its Graphical User Interface (GUI). *MorphData* takes advantage of IJM to automatically collect morphological features, working on any operating system in which ImageJ can work. The plugin is open-source and available online to the community. A straightforward architectural diagram is depicted in Figure 2.

**Figure 2.**
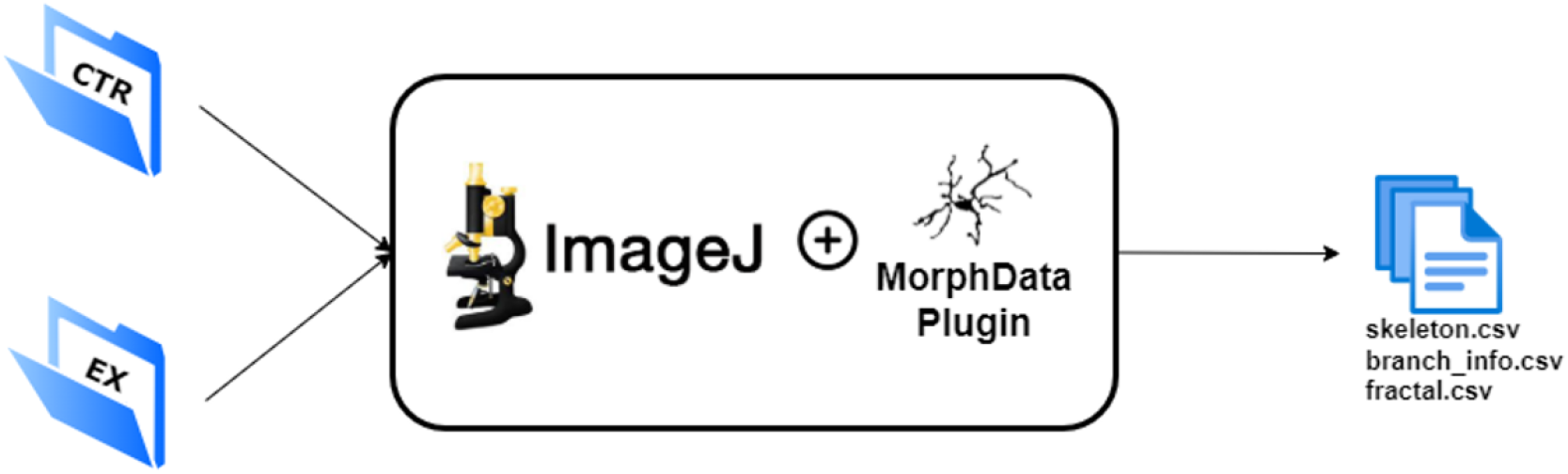
*MorphData*’s architectural diagram, receiving, as input, the root folders, and producing, as output, three csv files with the morphological features that characterize each single cell.

### 3.1 Computational requirements

The *MorphData* plugin requires basic computational resources. The experiments here described were carried out on a personal computer with an 8^th^ generation i7 CPU with 4 cores at 1.80GHz, 8GB of RAM, a SSD disk, and the Windows 10 operating system. ImageJ 1.53c, embedded in Fiji, has been set with 6989MB of maximum heap size.

*MorphData* runs in any operating system compatible with ImageJ (https://imagej.nih.gov/ij/index.html), which is available, as a downloadable application, for Windows, macOS, and Linux. The plugin is reliant on ImageJ (version 1.52t, or later) and the following ImageJ plugins:

i. *AnalyzeSkeleton (2D/3D)* (version 3.4.2, or later);
ii. *FracLac* (version 2015Sep090313a9330, or later).

ImageJ/Fiji requires a system with a Java 8, or later, virtual machine. *MorphData*’s post-processing script requires Python (version 3.7.10, or later) and the following modules:

- pandas (version 1.2.3, or later);
- tkinter (version 8.6, or later).

### 3.2 Installation

To install the *MorphData* plugin the user is required to download ImageJ and associated bundles with preinstalled plugins, such as Fiji, prior to installation (imagej.net/Fiji/Downloads). The user is then required to add *MorphData* as a new plugin to ImageJ:

i. Download the *Morph_Data*.*ijm* file from the code repository;
ii. Put the file in the plugins folder of ImageJ/Fiji itself.

The user can then start ImageJ and the *MorphData* plugin will be available at the Plugins tab. A detailed description on how to install ImageJ plugins can be found online at https://imagej.net/plugins.

*MorphData* also comes with a post-processing script, which users can run if necessary. To use this script users are required to have a Python environment installed. The easiest way to have such an environment is to download and install Anaconda, a popular open-source Python distribution platform (www.anaconda.com/products/individual). To run the post-processing script the user must open the Python console/prompt and execute the command “*python MorphData_PostProcessing*.*py*”.

### 3.3 Algorithm

Before detailing *MorphData*’s algorithm, it is important to clearly structure the obtained single cell images in the file system. Ideally, the user should create a structure such as the one depicted in Figure 3. To comply with the *MorphData* plugin, while the name of the folders at the two first levels is irrelevant, it is important to guarantee that the last two levels are entitled as “*Slice i*”, where *i* identifies different slices, and “*Image j*”, where *j* identifies different photomicrographs. Single cells should be placed inside the corresponding image folder, being entitled as “*Microgliak*.*tif*”, where *k* identifies each cell within the image folder.

**Figure 3.**
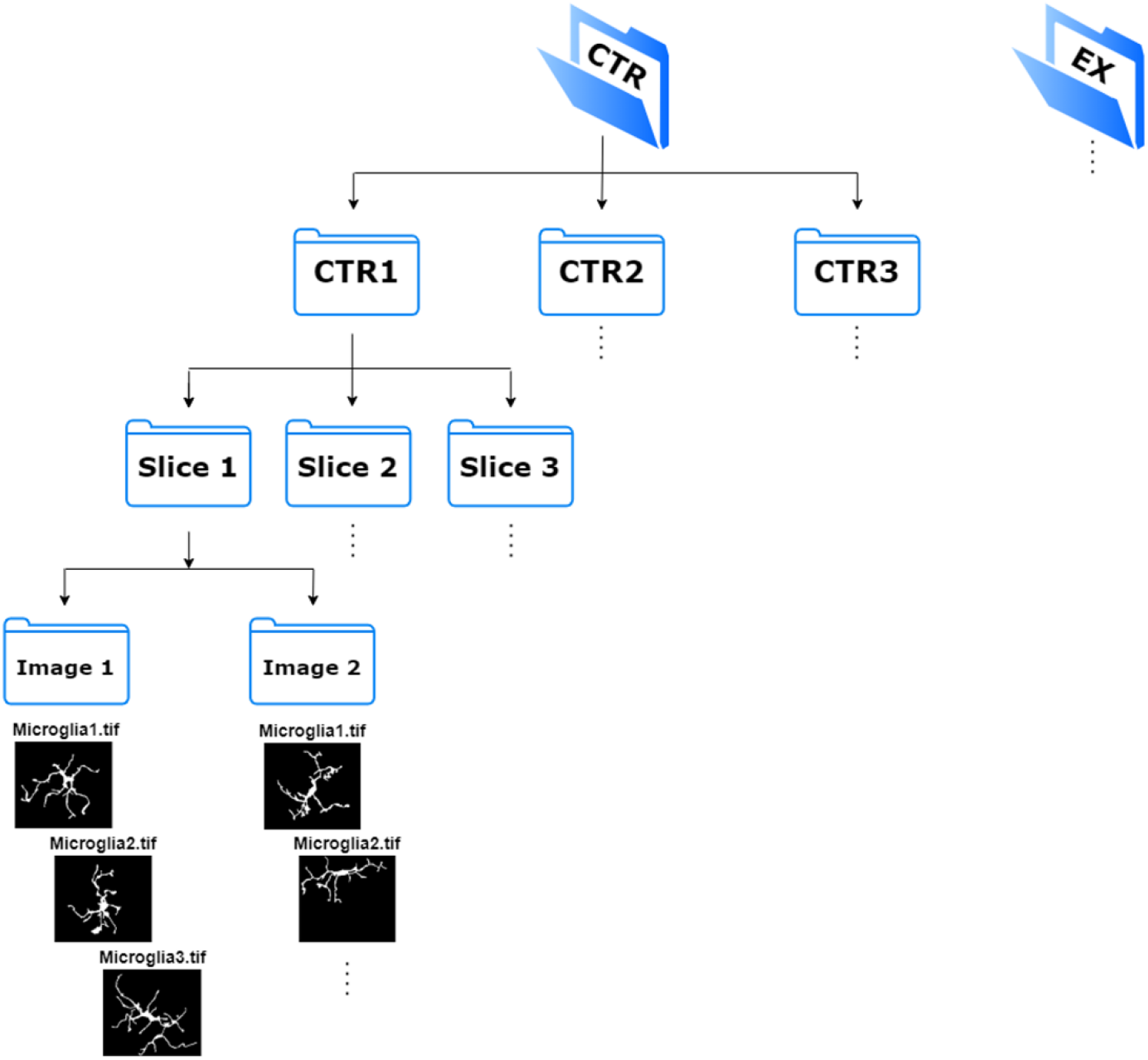
Recommended file system structure to store single cell images. The root folders, CTR and EX, hold the images of the corresponding experimental group.

The *MorphData* plugin, when executed, starts by asking the user to indicate the folder containing the single cell images (Figure 4a). Following the file structure defined in Figure 3, the user should indicate the *CTR* folder (the root folder). The plugin then creates auxiliary folders to store the collected data and automatically starts navigating the indicated folder looking for single cell images. Then, for each image, the algorithm is summarized as follows:

**Figure 4.**
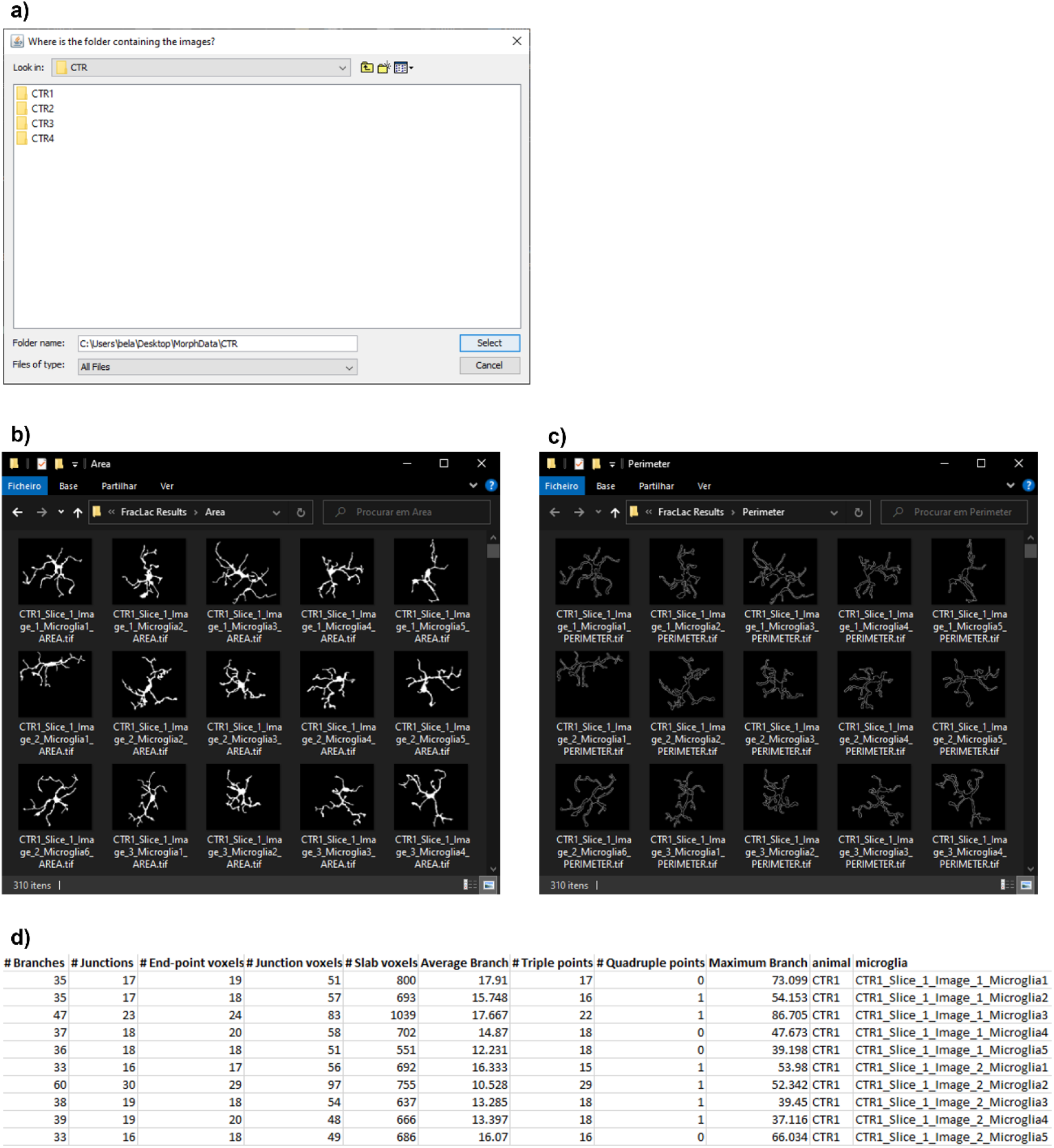
Execution and results of the *MorphData* plugin. **a)** *MorphData* dialog GUI asking the user where the single cell images are located. **b)** Shaped images, produced by *MorphData*, stored in the “*Area*” folder. **c)** Outlined images, produced by *MorphData*, stored in the “*Perimeter*” folder. Both shaped and outlined images are ready to be passed to the *FracLac* plugin for batch mode execution. **d)** A sample of the *skeleton_final_results*.*csv* file, produced by *MorphData*. This file contains 11 features relevant to cell ramification and cell identification.

i. To obtain skeletal features relevant to cell ramification:
  a. Open a single cell;
  b. Run the command “*Process* > *Binary* > *Skeletonize*” to create a skeletonized image;
  c. Run the *Analyze Skeleton (2D/3D)* plugin;
  d. Run the “*saveAs*” command to collect and store, in a csv file, skeletal data;
  e. Run the “*saveAs*” command to collect and store, in a csv file, branch information data.
ii. To obtain features relevant to cell complexity:
  a. Open a single cell;
  b. Run the “*saveAs*” command to store a shaped single cell, in TIFF format, in a folder entitled as “*Area*”;
  c. Run the “*Process* > *Binary* > *Outline’’* and “*saveAs*” commands to store an outlined single cell, in TIFF format, in a folder entitled as “*Perimeter*”.
iii. Repeat steps 1. and 2. for each single cell;
iv. At the end, the algorithm indicates the total number of analyzed cells.

Finally, contrary to the *Analyze Skeleton (2D/3D)* plugin, which is automatically executed by *MorphData*, the *FracLac* plugin cannot be directly executed from within another plugin. This limitation requires the user to manually execute the *FracLac* plugin itself after the *MorphData* plugin has finished. Fortunately, since the “*Area*” and the “*Perimeter*” folders, which were automatically created by *MorphData*, already contain all shaped and outlined cells (Figure 4b and 4c), the user can execute the *FracLac* plugin in batch mode. Hence, with a batch execution of this plugin, the user obtains fractal data for all cells almost immediately (avoiding the need to execute *FracLac* for each cell individually).

### 3.4 Post-processing script

Up to this point, all morphological data are now available, for all single cells, in multiple csv files in auxiliary “*results*” folders. In total, the *MorphData* plugin gathers 221 features (from skeleton to fractal ones), and some of them may be irrelevant to the characterization of microglial cells. Hence, the post-processing step is useful to join all data, cleaning irrelevant features, and performing a feature engineering process to create new ones, including the *cell_area, cell_perimeter, roughness*, and *cell_circularity*, among others.

Due to the potential high number of rows (cells) and columns (morphological features), an ImageJ plugin is unsuitable for the task, as it would eventually run out of memory. Hence, a Python script, entitled as *MorphData_PostProcessing*.*py*, was conceived and released as part of the *MorphData* plugin. This script requires a simple python environment to execute, again asking the user to indicate the location of the root folder. It will then automatically apply the post-processing procedures, creating three final files, containing the following 46 features:

i. **skeleton_final_results.csv:**
  - *# Branches, # Junctions, # End-point voxels, # Junction voxels, # Slab voxels, Average Branch Length, # Triple points, # Quadruple points, Maximum Branch Length, animal, microglia_id*.
ii. **branch_info_final_results.csv:**
  - *Skeleton ID, Branch length, V1 x, V1 y, V1 z, V2 x, V2 y, V2 z, Euclidean distance, running average length, average intensity (inner 3rd), average intensity, animal, microglia_id*.
iii. **fraclac_final_results.csv:**
  - *fractal_dimension, lacunarity, outline_mean_fg, density, span_ratio_major_minor, convex_hull_area, convex_hull_perimeter, convex_hull_circularity, diameter_bounding_circle, mean_radius, max_span_across_convex_hull, max_min_radii, shape_mean_fg, 1_pixel_side_micron, 1_pixel_area_micron_sq, cell_area, cell_perimeter, roughness, cell_circularity, animal, microglia_id*.

Figure 4d contains a graphical perspective of part of the content of the *skeleton_final_results*.*csv* file, which contains 11 features relevant for cell ramification. The remaining two files are similar, varying only on the quantified features. Sample input and output data are available at *MorphData*’s code repository.

## 4. Results and discussion

The performance of *MorphData* was evaluated based on the validity of the collected values and on the time it took to obtain the morphological data of all single cells of both experimental groups, when compared to a manual collection of such data.

Totally, in a computer with limited computing power, it took less than 14 minutes to collect 46 morphological features associated with 699 single cells of two experimental groups. In particular, 6.5 minutes were spent by the *MorphData* plugin, and its post-processing script, handling the CTR group. Of those, nearly 3 minutes were spent collecting skeleton data, 3.25 minutes by the *FracLac* plugin on batch mode, and 10 seconds by the post-processing script. On the other hand, 7.5 minutes were spent handling the EX group. Of those, 3.5 minutes were spent collecting skeleton data, 3.8 minutes by the *FracLac* plugin on batch mode, and 11 seconds by the post-processing script.

The same process was performed manually, by a skilled user of ImageJ, for a set of ten single cells of the CTR group. To ease the process, the same file system structure (as required by the *MorphData* plugin) was used. The goal was to mimic the processes that are automatically performed by *MorphData*, and manually collect 46 morphological features for the ten cells. The mean time to collect such morphological features was of 13 minutes per cell. Skeleton data were faster to collect (around 1.5 minutes), since the *AnalyzeSkeleton (2D/3D)* plugin only opens two results’ windows that the user can immediately save in two distinct files in the file system, in csv format, and then close the opened windows. However, fractal data were considerably harder to collect (around 11.5 minutes). On the one hand, for each cell, the *FracLac* plugin must be executed twice - one for a shaped cell and one for an outlined cell, which the user must prepare. On the other hand, for each execution, this plugin opens multiple results’ windows. The ones to keep opened are the “Hull and Circle Results” and the “Box Count Summary” windows. However, these windows are not user-friendly and, besides providing the user with an overwhelming amount of 173 features (most of them formulas and unwanted columns), it does not allow the user to copy only the desired features - the user must manually write each value of each desired feature to a csv, or excel, file. In fact, the process of selecting features from the *FracLac* plugin is extremely exhausting and error-prone. Finally, it is up to the user to calculate the value of non-existing features such as *cell_area, cell_perimeter, roughness*, and *cell_circularity*.

A couple more obstacles emerged with the manual process. First, the user was required to edit each stored file, for each cell, to identify the animal and the microglia of each row of data. Secondly, the user was required to copy the contents of each file to an overall file, aggregating the data for the experimental group. Since each cell is made of three files (two skeleton files and one fractal file) this would require the user to open and copy 699×3 files, which would, again, be a time-consuming task that would have to be performed after collecting all data.

Overall, the manual process for ten single cells of the CTR group took more than 2 hours to complete. On the other hand, the *MorphData* plugin and its post-processing script took less than 14 minutes to collect, process, and organize the morphological features of 699 cells. Assuming a mean value of 13 minutes per cell, the manual process to collect the morphological features of all cells would take 151 hours, which corresponds to almost 19 working days (8 hours/day) collecting data without stopping.

*MorphData* brings obvious advantages, mainly by significantly reducing the time it takes to collect morphological data. These values could be further reduced by a computer with higher computation power. In addition, the automation of the data collection process completely removes the risk of human error. It is worth mentioning that since *MorphData* is using well established plugins to collect morphological features, it produces the same exact results as when performing the data collection process manually. In fact, *MorphData*’s collected values were further compared and validated with multiple cells data that were manually collected by multiple people, without a single collection error.

## 5. Concluding Remarks

Morphological characterization of cells is highly relevant in the sciences field, and particularly in neurosciences. However, when performed manually, the process for obtaining morphological features from single cells is a demanding, repetitive, and laborious task; it is error-prone and can take several weeks to complete.

*MorphData* has already been used successfully in morphological characterization studies, where several thousands of single microglial cells, from multiple experimental groups, were analyzed and characterized. The benefits were considerable - several weeks of work were spared. Even though the plugin was optimized for microglial cells, it is likely to be performant for other glial cells, such as astrocytes and oligodendrocytes, and non-glial cells, such as neurons. Likewise, *MorphData* can also be used to automate the data extraction process of morphological features of *in vitro* cells.

It is an open-source plugin. Hence, new contributors, of all experience levels, are welcome. Contributions can be proposed using the pull request feature of GitHub or by opening a new issue (github.com/anabelacampos/MorphData/issues). These contributions can, among others, focus on the data extraction process, on *MorphData* performance over different cell types other than microglial cells, improve the documentation, or be made of constructive feedback and suggestions.

Overall, *MorphData* significantly reduces the time taken to collect morphological data from microglial cells (from weeks to minutes), which can then be used to study, understand, and characterize microglia behavior in the brain of human patients or of animal models of neurological and psychiatric diseases.

## Acknowledgements

This work is supported by *Fundação para a Ciência e a Tecnologia* (FCT) (PTDC/NEU-NMC/3648/2014) and COMPETE-FEDER (POCI-01-0145-FEDER-016818). ABC is supported by a doctoral fellowship from FCT (PD/BD/127828/2016).

## Authors Contributions

ABC: Conceptualization; Methodology; Validation; Formal analysis; Investigation; Data Curation; Writing - Original Draft; Writing - Review & Editing; Visualization. SDS: Conceptualization; Resources; Writing - Review & Editing; Supervision. AFA: Conceptualization; Resources; Writing - Review & Editing; Supervision; Funding acquisition. PM: Conceptualization; Resources; Writing - Review & Editing; Supervision; Funding acquisition. BF: Methodology; Software; Validation; Formal analysis; Data Curation; Writing - Review & Editing; Visualization

## Data Accessibility

Sample input and output data are provided to the public for educational and academic research purposes, being freely available at *MorphData*’s code repository.

## Abbreviations

CTR: control
EX: experimental group
GUI: Graphical User Interface
IJM: ImageJ Macro language
PBS: Phosphate-Buffered Saline
RT: room temperature

